# Learning feedback loops in transcriptome network during cell fate transition

**DOI:** 10.1101/2020.06.24.169490

**Authors:** Lingyun Xiong, William A. Schoenberg, Jeremy D. Swartz

**Affiliations:** Department of Integrative Biology and Physiology, University of California, Los Angeles, CA USA; Institute for Quantitative and Computational Biosciences, University of California, Los Angeles, CA USA; Department of Geography, University of Bergen, Norway; isee systems inc. Lebanon NH, USA; Media Studies Program, University of Oregon, USA

**Keywords:** nonlinear feedback, network dynamics, machine learning, ordinary differential equation, cell fate transition

## Abstract

Embryonic development is characterized by series of cell fate transitions, during which non-linear feedback of signaling events and network dynamics in the transcriptome are instrumental to determine differentiation outcomes. Changes in selected genes or pathways have been described extensively, but a system-level understanding of gene regulatory network regulation is still lacking. Leveraging time-series data from single-cell transcriptome profiling of developing mouse embryos, we collapsed dynamic gene expressions into changes in 14 core biological processes, and constructed an ordinary differential equation-based machine learning model of the transcriptomic network to capture its feedback structure. We evaluated the polarity and magnitude of direct pairwise causal relationships between biological processes and identified higher-order feedback loops that dominate system behavior. Despite heterogeneous expressions at the gene level, we find that the transcriptome at the macroscale level has a feedback structure that is intrinsically stable and robust. We show the pivotal role of intracellular signaling in driving systemic changes and uncover the importance of homeostatic process and establishment of localization in regulating network dynamics. Local regulatory structures represent domain-specific regulations that are essential to cell fate transition, especially lipid metabolic process. Together, this study provides a holistic picture of transcriptomic network dynamics during mouse organogenesis, offering insight into key aspects of information flow in the transcriptome that control cell fate transition.

**Conflict of interest statement:** The authors declare no potential conflicts of interest.

**Funding:** The authors received no funding for this study.

**Author contributions:** Conceptualization: LX; Methodology: LX, WAS, and JDS; Data Curation: LX; Investigation: LX and WAS; Formal Analysis: LX and WAS; Visualization: WAS and LX;

Writing – review & editing: LX and WAS.

## 1 Introduction

Cell fate transition is the central theme of embryonic development: embryonic stem cells diversify into intermediate progenitors that will differentiate into diverse cell lineages of specialized form and function in all organs (Barresi & Gilbert, 2024). This is a complex yet well-coordinated process, effected by the spatial and temporal dynamics of gene expression (Arnold & Robertson, 2009; Sladitschek et al., 2020; Young, 2011). Cell identity can be summarized by the collective state of the transcriptome (Cao et al., 2019; Cheng et al., 2019; Macarthur, Ma’ayan, & Lemischka, 2009; Pijuan-Sala et al., 2019). Recent years has seen tremendous effort in probing transcriptome-wide gene expression during mouse organogenesis at single-cell resolution. This offers a unique opportunity to look beyond individual genes and pathways (Tam & Loebel, 2007), to model the dynamics of gene regulatory network as a whole (Davidson et al., 2002; Lu et al., 2009). In particular, Pijuan-Sala et al. sequentially sampled developing mouse embryos every 6 hours between 6.5 to 8.5 days post-fertilization (inclusive) and performed single-cell RNA sequencing to construct the cell atlas during early mouse organogenesis. Utilizing this comprehensive dataset, we aim to establish a formal description of the transcriptome dynamics that underlies cell fate transition.

With more than 20,000 genes annotated in the mouse genome, the gene regulatory network is high-dimensional in nature. So far, analyses of biological networks have mainly focused on sub-network, such as transcriptional regulatory networks (Luscombe et al., 2004; Neph et al., 2012), protein-protein interaction networks (Fraser, Hirsh, Steinmetz, Scharfe, & Feldman, 2002; Han et al., 2004), and metabolic networks (Fondi & Liò, 2015; Jeong, Tombor, Albert, Oltvai, & Barabási, 2000). Gene Ontology (GO) is a literature-curated reference database that annotates genes’ functions in the hierarchy of cellular subsystems (Bult et al., 2019; Ma et al., 2018; The Gene Ontology Consortium, 2017), which can be beneficial to reduce the dimensionality of the network while capturing its entirety. Taking this knowledge about genes and cellular subsystems as an anchor point, we can model the gene regulatory network in a coarse-grain manner, where individual genes can be collapsed into 14 core biological processes as annotated by the Mouse Genome Institute.

Formalism of modeling network dynamics in biological systems is severely constrained by the presence of nonlinear and feedback-rich interactions among network components (L. Xiong & Garfinkel, 2022; Zou, Donner, Marwan, Donges, & Kurths, 2019). Existing paradigms draw inference either on pre-specified (sub-)network structures (Harush & Barzel, 2017) or using probabilistic techniques without identifying the underlying causal structure (Casadiego, Nitzan, Hallerberg, & Timme, 2017; Groß et al., 2019). Here, we take a data-driven approach with a mathematical backbone to model the system dynamics in terms of network structure (Aibar et al., 2017; Fujii, Takeishi, Hojo, Inaba, & Kawahara, 2020; Peixoto & Rosvall, 2017; Tu, Kudlicki, Rowicka, & McKnight, 2005). We agnostically train a causal network model that competently reproduces network dynamics. We then instrument the generated model to infer the nature of interactions from the causal structure of the mathematical model.

In this study, we describe the macroscale dynamics of the transcriptome during mouse organogenesis using a causal machine learning model based on ordinary differential equations. We developed an analytic framework for modeling the high-dimensional gene regulatory network at the macroscale, which algorithmically generates a network model to explain system behavior. We find that the feedback structure of the macroscale transcriptome network is intrinsically stable and robust. Among the 14 core biological processes, *signaling* plays a pivotal role in regulating system dynamics during mouse organogenesis, and *homeostatic process* and *establishment of localization* exhibit secondary effect. Tertiary regulatory processes include *system development* and *cell differentiation*. Interestingly, *lipid metabolic process* exerts significant impact on the network dynamics by negative feedback on multiple core biological processes, which is potentially entangled in a bistable relationship with *signaling* in facilitating cell fate transition.

## 2 Results

### 2.1 A coarse-grain representation of the gene regulatory network

In Gene ontology (GO), genes are annotated for its functions in diverse molecular/biochemical processes, such as DNA repair and cell growth (The Gene Ontology Consortium, 2017). Fine or coarse, the annotations are related in a largely hierarchical directed acyclic graph (DAG). Child-parent relationships in the DAG link lower-level subsystem-oriented functions to those pertaining to upper-level broad categories. We traced the paths in the DAG and collapsed the lower-level terms to the top-level ones, condensing the gene regulatory network into a 14-node representation (i.e., the macroscale network; Fig 1A and 3; *Methods*). Each node encapsulates genes involved in a distinct functional domain of the biological system (Table 1; Table S1 and S2) (Bult et al., 2019).

**Table 1:**
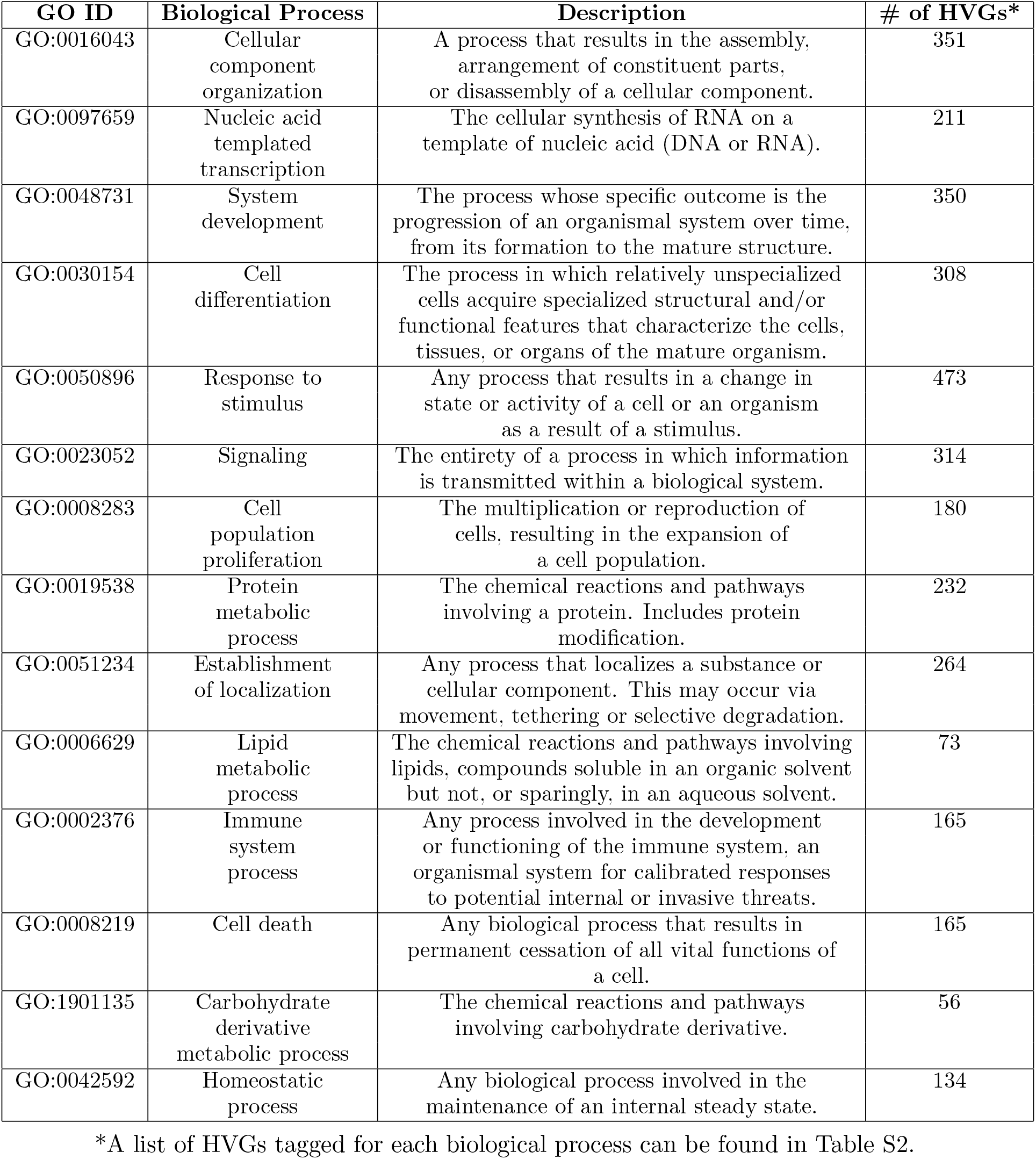
Summary of the 14 core biological processes based on Gene Ontology annotations.

| GO ID | Biological Process | Description | # of HVGs* |
| --- | --- | --- | --- |
| GO:0016043 | Cellular component organization | A process that results in the assembly, arrangement of constituent parts, or disassembly of a cellular component. | 351 |
| GO:0097659 | Nucleic acid templated transcription | The cellular synthesis of RNA on a template of nucleic acid (DNA or RNA). | 211 |
| GO:0048731 | System development | The process whose specific outcome is the progression of an organismal system over time, from its formation to the mature structure. | 350 |
| GO:0030154 | Cell differentiation | The process in which relatively unspecialized cells acquire specialized structural and/or functional features that characterize the cells, tissues, or organs of the mature organism. | 308 |
| GO:0050896 | Response to stimulus | Any process that results in a change in state or activity of a cell or an organism as a result of a stimulus. | 473 |
| GO:0023052 | Signaling | The entirety of a process in which information is transmitted within a biological system. | 314 |
| GO:0008283 | Cell population proliferation | The multiplication or reproduction of cells, resulting in the expansion of a cell population. | 180 |
| GO:0019538 | Protein metabolic process | The chemical reactions and pathways involving a protein. Includes protein modification. | 232 |
| GO:0051234 | Establishment of localization | Any process that localizes a substance or cellular component. This may occur via movement, tethering or selective degradation. | 264 |
| GO:0006629 | Lipid metabolic process | The chemical reactions and pathways involving lipids, compounds soluble in an organic solvent but not, or sparingly, in an aqueous solvent. | 73 |
| GO:0002376 | Immune system process | Any process involved in the development or functioning of the immune system, an organismal system for calibrated responses to potential internal or invasive threats. | 165 |
| GO:0008219 | Cell death | Any biological process that results in permanent cessation of all vital functions of a cell. | 165 |
| GO:1901135 | Carbohydrate derivative metabolic process | The chemical reactions and pathways involving carbohydrate derivative. | 56 |
| GO:0042592 | Homeostatic process | Any biological process involved in the maintenance of an internal steady state. | 134 |

\*A list of HVGs tagged for each biological process can be found in Table S2.

**Figure 1.**
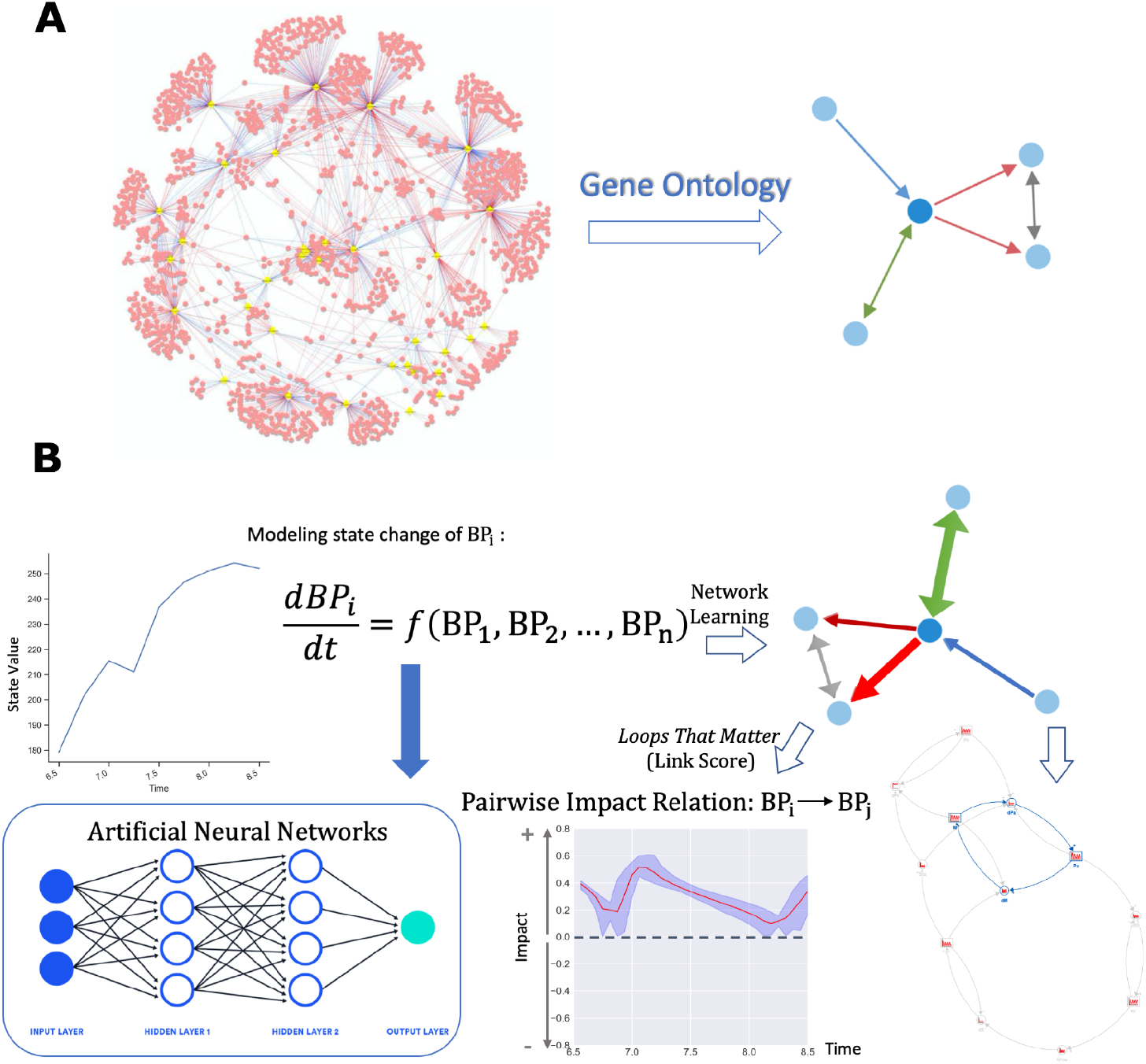
Graphical summary of study methodology. (A) Leveraging Gene Ontology annotations, we can aggregate individual genes in the entire gene regulatory network into a 14-node network at the macroscale level. Each node in the coarse-grain representation stands for a top-level biological process that are essential for cellular function. (B) To agnostically construct a causal model of the macroscale network that recapitulates the observed system dynamics, we used the Feedback System Neural Network (FSNN) method (Schoenberg, 2019) based on the aggregate gene expression levels in each core biological process. In brief, state change of each core biological process is modeled by a differential equation, whose form is captured by an artificial neural network. The machine learning model was trained and tested. The generated model can be analyzed mathematically to infer direct interactions among the core biological processes, in terms of both pairwise causal relationships and feedback loops that dominate system behavior.

Single-cell RNA sequencing data of development mouse embryos generated by Pijuan-Sala et al. was retrieved and processed according to authors’ instructions. This provides a matrix of normalized expression values of genes in respective cells. Cells are marked for specific stage they are sampled (E6.5-8.5, sampled every 6 hours, 9 time points in total). We selected cells annotated as endoderm lineage and we modeled variance of gene expression levels amongst these cells. We identified 982 highly variable genes (HVGs; Fig 2A; *Methods*), many of which are master regulators of embryo development (e.g., CDX2, OCT3/4, WNT5/6 and BMP2/4; Table S2) and biomarkers for gut organogenesis (e.g., Krt8, Krt18, Epcam, and Apela; Table S2). There are two converging trajectories of differentiation during gut organogenesis (Pijuan-Sala et al., 2019), one from anterior primitive streak to definitive endoderm to gut (T1), and the other from visceral endoderm to gut (T2). We thus split the cells into two, corresponding to the distinct but related cell populations. Next, we conducted a two-stage data aggregation scheme to group gene expression levels to represent the state value (i.e. gross activity) of each biological process (*Methods*). As a result, we have generated 14 time-series in every set (two trajectories, 4 sets each; Fig 2C; *Methods*). These sequential aggregate gene expression levels constitute the empirical basis of the downstream modeling and analysis.

**Figure 2.**
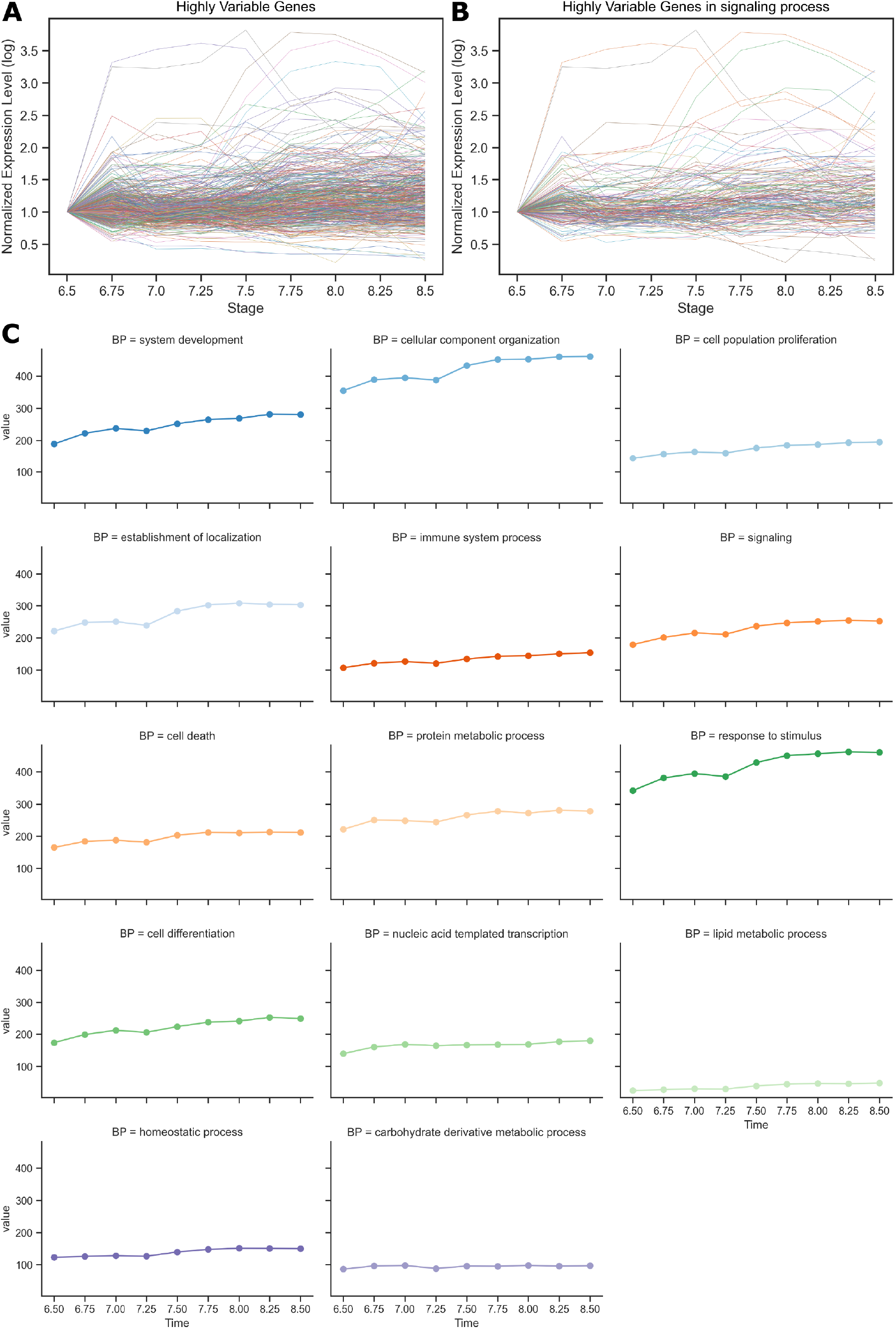
Aggregation of gene expression values to represent the gross activity of the 14 core biological processes (T1). (A) Average gene expression levels of highly variable genes during mouse gut organogenesis (E6.5-8.5, sampled every 6 hours). Expression values were normalized to the initial time point (E6.5) for visualization. (B) Average gene expression levels of highly variable genes tagged for *signaling*. (C) Aggregate gene expression levels (i.e. state values) of the 14 core biological processes.

### 2.2 A causal machine learning model of the macroscale network

Based on the state values of the 14 biological processes at the 9 sequential time points, we constructed a causal model of the macroscale network using Feedback System Neural Networks (FSNN; Fig 1B; *Methods*) (Schoenberg, 2019). For the two differentiation trajectories mentioned above, we generated two unique models: T1 and T2 (Fig 3). The trained models reproduced the test datasets, with an average mean squared error (R2) of 93% for T1 (range: [44.9%, 99.3%]) and 79% for T2 (range: [43.3%, 97.3%]), respectively (Fig 3A and 3C; Table S3).

**Figure 3.**
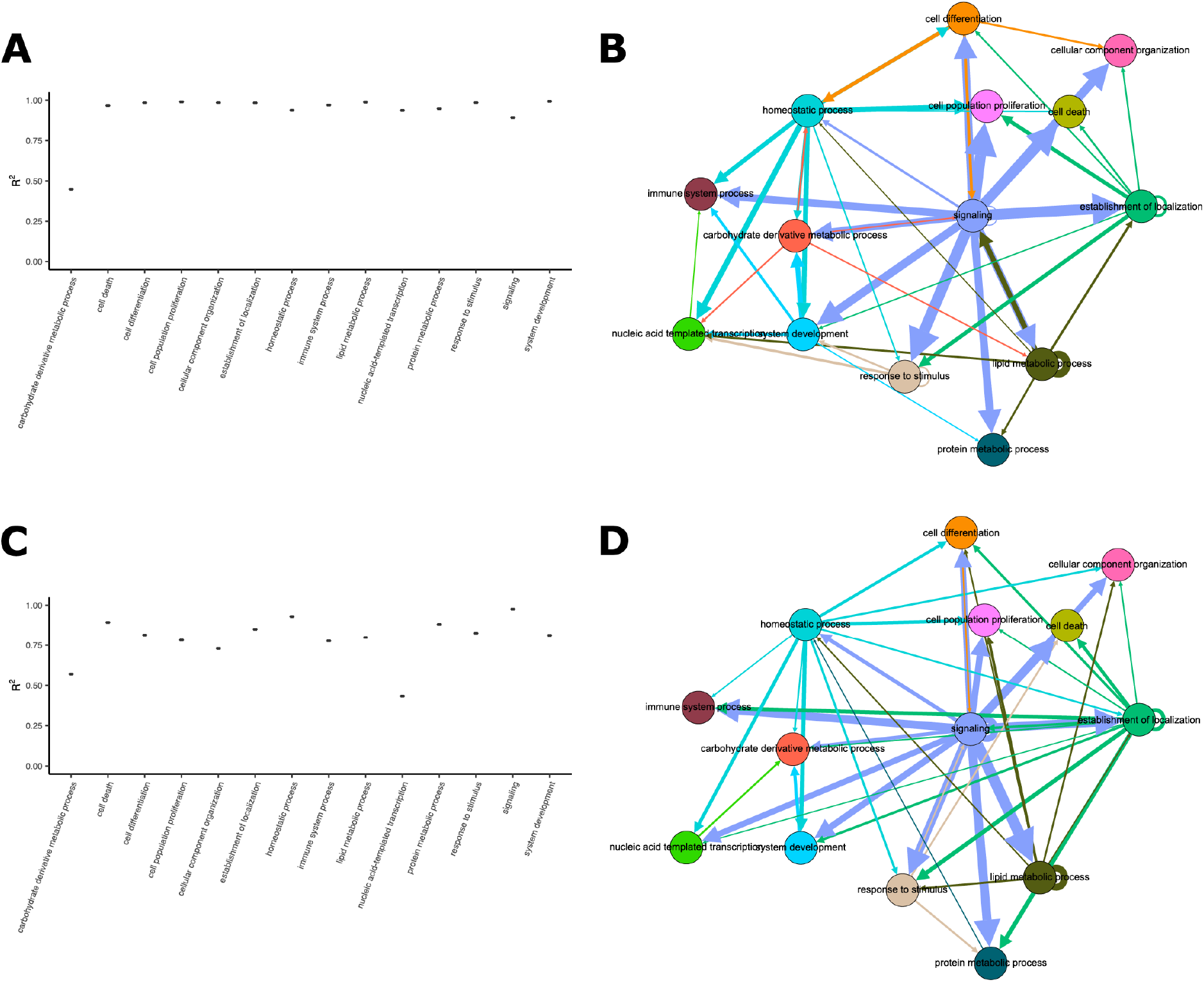
Two causal models of macroscale gene regulatory network during mouse gut organogenesis. (A, B) Trajectory 1 (T1): model is trained and tested on data drawn from cells annotated as anterior primitive streak, definitive endoderm and gut. (C, D) Trajectory 2 (T2): model is trained and tested on data drawn from cells annotated as visceral endoderm and gut. (A, C) Outcome of model parameterization, shown as mean square error (R2) measured between model prediction and test data for each biological process. (B, D) Static view of the network feedback structures. Nodes represent the 14 core biological processes. Directed edges indicate the presence of causal relationship from the state change in source node to the state change in target node. Edge width is proportional to the cumulative impact over the entire simulation period. Note that the arrow does not indicate the polarity of feedback (positive/negative). Only the strongest 49 edges (the upper quartile of all 196 possible directed connections) are included for visualization.

We analyzed the structure of the ordinary differential equations, to determine direct pairwise causal relationships between network components as well as dominant higher-order interactions (*Methods*). The link score metric was used to quantify the relative contribution of the state change in source node to the state change in target node (*Methods*). The link score measures both the polarity and magnitude of the causal relationship of interest. If the link score is positive (i.e. the signs of the first derivative of the source and target nodes are consistent), the feedback relationship is reinforcing; if negative (i.e. the signs of the first derivative of the source and target nodes are opposite), the feedback relationship is inhibitory. Ranging from 0 to 1, the magnitude of the link score reflects the strength of the causal relationship, which is calculated as the fraction of state change in target node accounted for by the state change in source node. Key higher-order feedback interactions were assessed and ranked via feedback loop dominance analysis (Schoenberg, Davidsen, & Eberlein, 2020; *Methods*). Static visualization of the network feedback structure demonstrates the cumulative impact of each node on its target nodes over the entire simulation period (Fig 3B and 3D). A dynamic view is offered by stacking frames of the instantaneous impact relations over time (Video S1 and S2).

Biological systems are known to harbor scale-free networks. Indeed, the degree of nodes in our causal models approximately follows a power-law distribution, with an average degree of 3.5 and an average path length of 1.8 (Fig S3). Despite the heterogeneity in gene expressions during embryo development (Fig 2A and 2B), we observe that the macroscale network is intrinsically stable and robust, as seen in the concordant network structure for both trajectories (Fig 3B and 3D; Video S1 and S2).

In the following sections, we present three commonalities in the network feedback structure: the primary role of *signaling* in regulating network dynamics during cell fate transition, the secondary roles of *homeostatic process* and *establishment of localization*, and localized impact structure that are crucial for cell fate transition.

### 2.3 The pivotal role of *signaling* in driving systemic changes

Central in the network models is *signaling*, whose state change induces state changes across the entire network (Purple edges; Fig 3B and 3D). Embodying “the entirety of a process of information transfer within a biological system” (The Gene Ontology Consortium, 2017; Table 1; Table S1), *signaling* plays a pivotal role of regulating network dynamics during mouse organogenesis, during which signal transduction has been regarded quintessential (Basson, 2012). Constituent genes in signaling mainly encode transcription factors (e.g., OCT3/4, GATA4, Sox family proteins, and T-box family proteins; Table S2) and key factors in signal transduction pathways (e.g., WNT pathway, TGF-β pathway, FGF pathway, and heat shock response; Table S2).

State change in *signaling* induces and reinforces the state changes in all 14 biological processes, including itself (Fig 4A; Fig S1 and S2). Its impact accounts for up to 70% of the state changes in its target nodes, including *system development* and *cell differentiation*, the canonical players in mouse embryo development (Bult et al., 2019; The Gene Ontology Consortium, 2017; Table 1; Table S1). Furthermore, among all the higher-order interactions inferred from the causal models, positive feedback of signaling on itself (loop R1) is one of the most prominent feedback loops that explain system behavior. It accounts for 2.24% and 2.12% of the total of system behavior in T1 and T2, respectively (Fig 3 and 5; Table S4 and S5). This positive feedback loop is necessary for the system to be bistable or multistable during cell fate transition (Ferrell, 2012). This feature of network topology reflects fundamental design principles of biological computation in the system.

**Figure 4.**
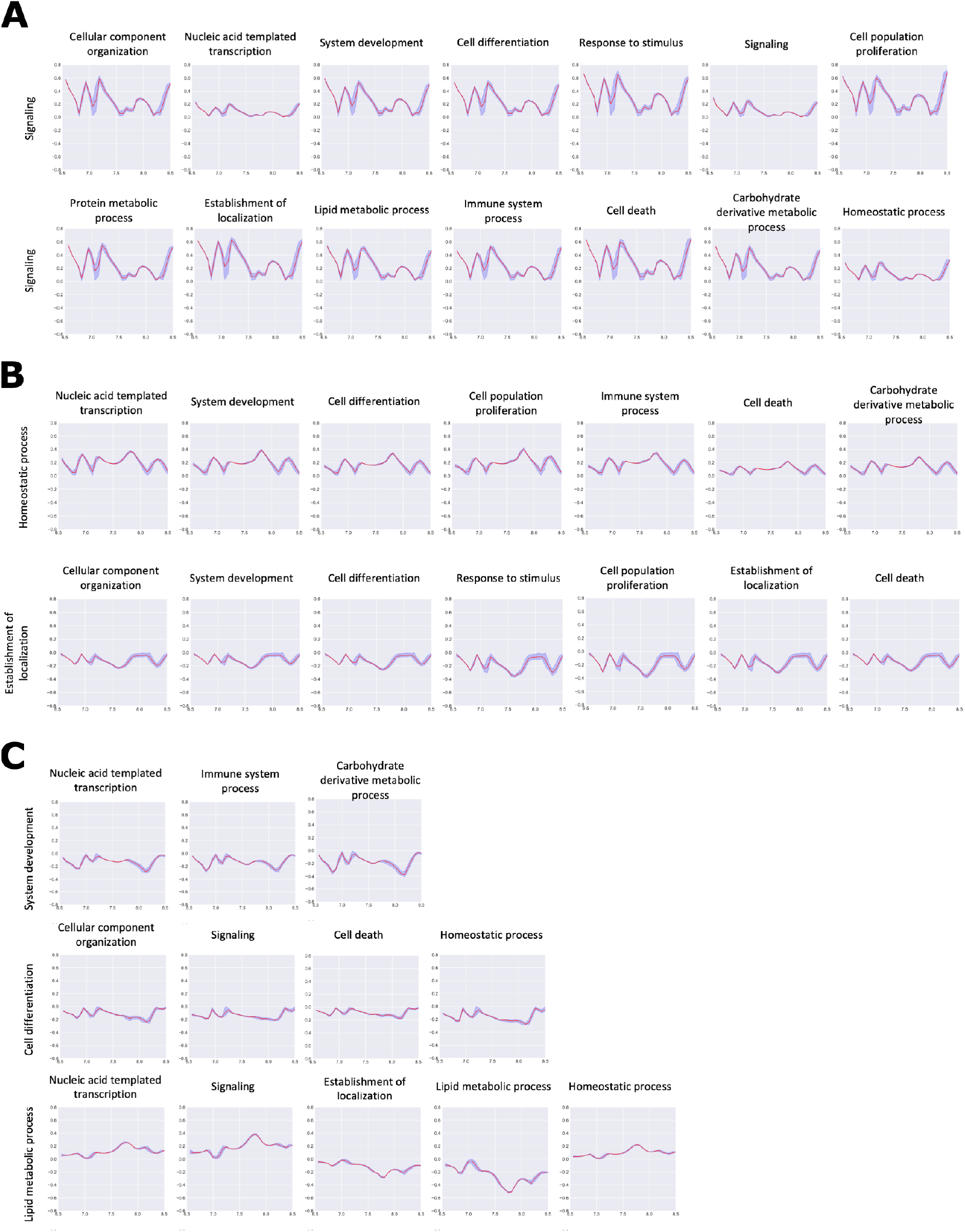
Pairwise causal relationships can be inferred from link score measures (T1). (A) Impact of state change in *signaling* on the state changes in all 14 nodes of the macroscale network during the entire simulation period is measured using link score (*signaling* → target node of interest). (B) Impact of state changes in *homeostatic process* and *establishment of localization* on the state changes in half of the macroscale network. (C) Example of localized pairwise causal relationships directed from *system development, cell differentiation* and *lipid metabolic process*. See Fig S1 and S2 for additional inferred pairwise relationship.

**Figure 5.**
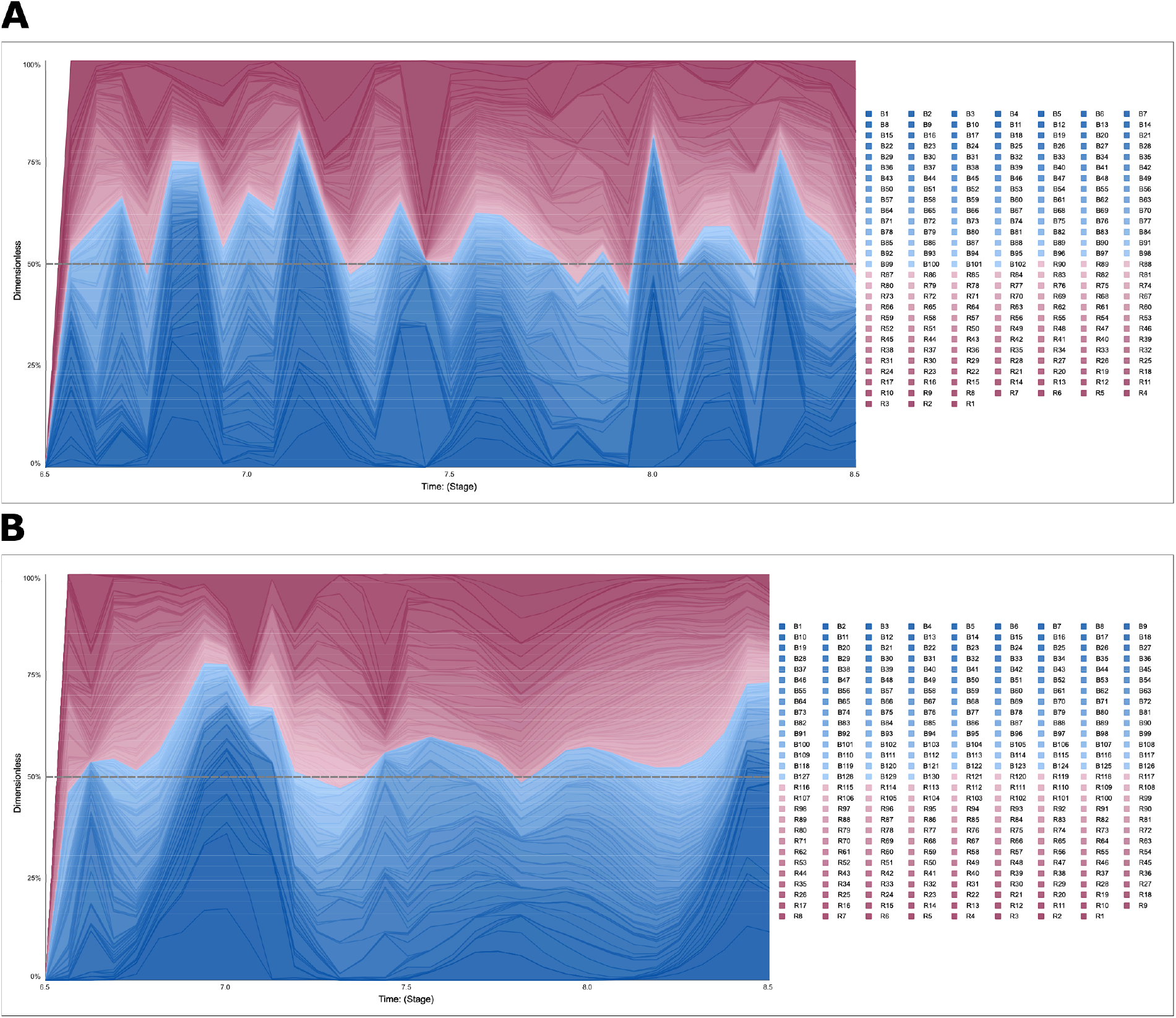
Inferred higher-order interactions in the causal models. Feedback loop dominance analysis of the two causal models (A: T1; B: T2) measures and ranks the impact of individual feedback loops on system dynamic behavior, shown here as the proportion of variance explained by each feedback loops during the simulation period (horizontal axis). Influence measures of individual feedback loops are shaded by two color gradients (red: positive/reinforcing feedback [even-numbered negative link scores in the loop]; blue: negative/inhibitory feedback [odd-numbered negative link scores in the loop]). The higher-ranked feedback loops are shaded in darker hues and arranged towards the extremes of vertical axis. Examples of the feedback loops presented: B1 – self-inhibitory feedback of *lipid-metabolic process* ; R1– self-reinforcing feedback of *signaling* ; B2 – self-inhibitory feedback of *establishment of localization*; B4 – negative feedback loop from *cellular component organization* → *signaling* → *cell differentiation* → *homeostatic process* → *system development* → *carbohydrate derivative metabolic process* → *nucleic acid templated transcription* → *cellular component organization* (Table S4 and S5).

### 2.4 Secondary roles of *homeostatic process* and *establishment of localization*

*Homeostatic process* and *establishment of localization* regulate dynamic changes in half the network (blue and green edges; Fig 3B and 3D). *Homeostatic process* refers to “any biological process involved in the maintenance of an internal steady state”(Bult et al., 2019; The Gene Ontology Consortium, 2017; Table 1; Table S1). Upon internal or external stimulus, products of constituent genes in *homeostatic process* issue instructions to specific cellular subsystems to restore system behavior, for example, to reorganize cytoskeleton and to switch between anabolic and catabolic processes in metabolism (L. I. Xiong & Garfinkel, 2023). Akin to *signaling*, it is in essence an ontological regulator process, which is critical for system regulation and control (Ferrell, 2016). Notably, state change in *homeostatic process* reinforces state changes in *system development* and *cell differentiation*, accounting for up to 40% of state changes (Fig 4B; Fig S1 and S2).

On the other hand, *establishment of localization* is an effector process, which executes mobility commands “via movement, tethering and selective degradation of molecules or cellular components”(Bult et al., 2019; The Gene Ontology Consortium, 2017; Table 1; Table S1). Its state change negatively regulates state changes in system development and cell differentiation, as well as itself (Fig 4B; Fig S1 and S2). The self-inhibitory feedback of establishment of localization (loop B2), like the self-reinforcing feedback of signaling (loop R1), plays an important role in the overall feedback loop structure: explaining 1.87% and 2.77% of the total system behavior in T1 and T2, respectively (Fig 3 and 5; Table S4 and S5). This indicates the important role of effector processes in regulating network dynamics.

### 2.5 Instances of localized impact structure

Localized interactions are also noted in the two network models, especially the two aforementioned canonical players in mouse embryo development: *system development* and *cell differentiation* (cyan and orange edges; Fig 3B and 3D; Table 1; Table S1). Markedly, state changes in *cell differentiation* negatively regulate state changes in *signaling* and *homeostatic process* (Fig 4C; Fig S1 and S2). *System development* negatively regulates *nucleic acid transcription*, consistent with previous reports (Pan, Li, Zhou, Zheng, & Pei, 2006; Papatsenko, Waghray, & Lemischka, 2018). Intriguingly, *system development* also influences on *carbohydrate derivative metabolic process* (with a magnitude up to 40%).

Strikingly, we observed pronounced negative feedback of *lipid metabolic process* on itself (loop B1) in both trajectories (brown edges; Fig 3B and 3D). State change in *lipid metabolic process* reinforces state changes in *signaling* while it counteracts state changes in *establishment of localization*, the primary and secondary hubs of the macroscale network (Fig 4C; Fig S1 and S2). Moreover, the self-inhibitory feedback of *lipid metabolic process* is if not more, at least as explanatory of the network dynamics as the self-reinforcement of signaling (loop R1): explaining 3.15% and 3.07% of the total system behavior in T1 and T2, respectively (Fig 3 and 5; Table S4 and S5). It is noted that loop B1 persistently precedes loop R1 in time (Fig 5). Considering the mutual pairwise causal relationships of *signaling* (Fig 4A) and *lipid metabolic process* (Fig 4C), the two functional domains are plausibly entangled in a relationship resembling a bistable switch (Fig 6). This relationship comprises a recurring scene of information flow during cell fate transition, echoing recent development on understanding the function of peroxisome and PPAR proteins (Ahmadian et al., 2013; Liu et al., 2018; Lodhi & Semenkovich, 2014). This represents an axis in regulating network dynamics during mouse organogenesis that warrants further investigation.

**Figure 6.**
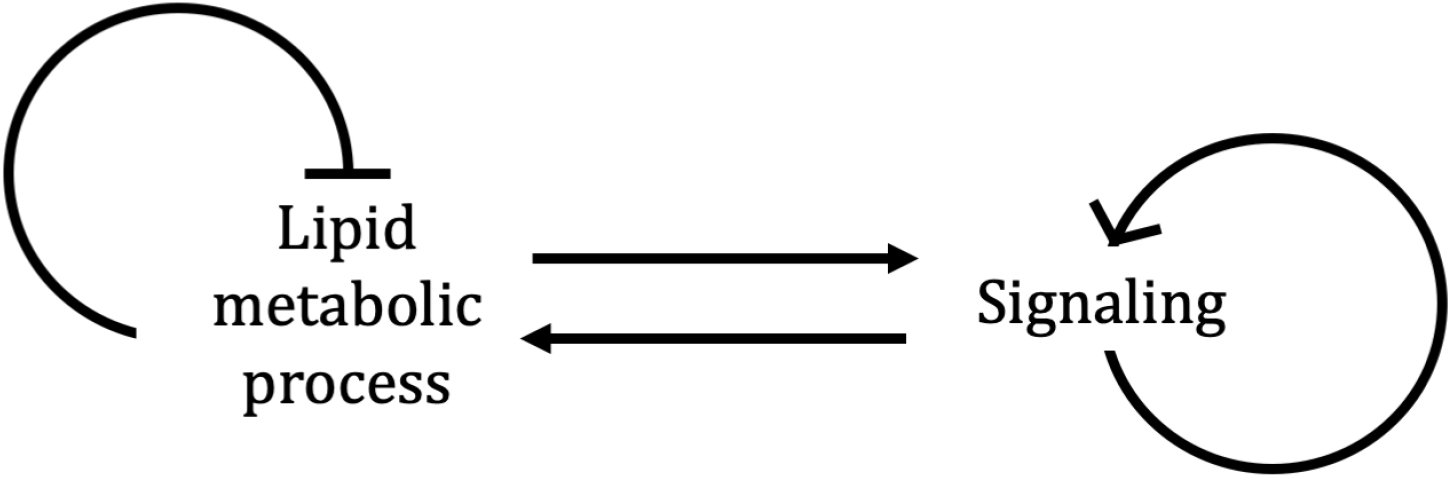
Hypothesized bistable feedback relationship between *lipid metabolic process* and *signaling*.

In addition, incidental impact of *carbohydrate derivative metabolic process* on *signaling, homeostatic process, nucleic acid templated transcription*, and *lipid metabolic process* is also noted (Fig S1 and S2). Metabolic state of the cell reflects the availability of nutrients and cellular capacity to use them effectively (Kang et al., 2024; Merkuri, Rothstein, & Simoes-Costa, 2024). Metabolic activity thus fine-tunes the signaling network in the cell (Causton, 2020; Gomes & Blenis, 2015; Wellen & Thompson, 2012). Together, this reveals the essential role of metabolism, as an effector of mass and energy flow, in the network feedback structure governing cell fate transition. Again, this emphasizes the interactions between regulator and effector processes in controlling network dynamics during organogenesis.

## 3 Discussion

Transcriptome dynamics underlie cell fate transition during embryo development. Besides the actions of master regulators and biomarkers, it is increasingly clear that the collective state of the transcription network reflects and effects cell identity. In this study, we provide a formal description of macroscale transcriptome network dynamics during mouse organogenesis by constructing a causal machine learning model and delineating the feedback structure embedded in the network topology. Our analysis characterizes key aspects of information flow across the network that are known to induce changes and uncovers features that have not been acknowledged in full. We point out new axes of inter-relationships between ontological regulator and effector processes, especially between *lipid metabolic process* and *signaling*. These interactions plausibly control information flow during mouse organogenesis, which warrants further investigation. Encompassing the entire transcriptome, we have thus integrated mass, energy, and information transfer within the same analytical framework.

The causal models we have generated for the macroscale gene regulatory network are time-dependent nonlinear feedback systems that is biologically interpretable, offering a dynamic view of the transcriptome during mouse organogenesis. The bird’s-eye view offers a vantage point for us to examine the large-scale organization of the system during a phase of drastic transformation at the organismal level. During this process, we observed that the macroscale network is highly organized and the information flow is intrinsically robust. This regularity prompts us to postulate that the macroscale feedback structure epitomizes the origin of stability of the system during its natural progression. If true, it would be of interest to compare our causal models to those generated from biological systems under pathological conditions, such as cancer (Rockne et al., 2020; Rozenblatt-Rosen et al., 2020). Meanwhile, this macroscale representation inevitably overshadows granularity that is required for experimental/clinical actions. Nevertheless, given the largely hierarchical organization of GO annotations, it is feasible to aggregate at the level of concrete subsystems, such as different pathways involved in DNA damage response, instead of the top-level biological processes. This allows for a mesoscale representation of the network, which is currently an active area of research (Goodsell, Olson, & Forli, 2020).

Due to pervasive nonlinear interactions and combinatorial circularity in feedbacks, network dynamics of biological systems remains a challenge to investigate (Kurz et al., 2017; Shin, Venturelli, & Zavala, 2019). In this article, we have inferred network feedback structure by constructing a causal model with no prior specification. Building upon network learning algorithm and numerical methods, this analytic framework not only offers means to visualize network dynamics, but also pinpoints essential characteristics of the network topology that underlie systemic dynamics. Many real-world networks share properties with gene regulatory network. It is conceivable that this analytic framework is readily applicable to these fields.

## 4 Methods

### 4.1 Data processing

Single-cell RNA-seq data from developing mouse embryos (C57BL/6 genetic background) over a time-course of 48 hours (embryonic day E6.5-E8.5) was generated and curated by Pijuan-Sala et al., (Pijuan-Sala et al., 2019). Raw mapped read counts covering 29,452 genes across the genome in 116,312 cells were downloaded from https://github.com/MarioniLab/EmbryoTimecourse2018. We processed the data following authors’ instructions, including doublet removal, reads normalization, and batch correction. After initial processing, cells of endoderm lineage were selected by clustering information available in meta-data (cluster 9 [excluding sub-cluster 1 & 3] and cluster 11). Cells were split into two developmental trajectories based on cell type annotations: trajectory 1 includes cells annotated as anterior primitive streak, definitive endoderm, and gut; trajectory 2 includes cells annotated as visceral endoderm and gut. Due to low sampling at early time points in trajectory 1, data imputation was performed to reach a balanced sample sizes in both trajectories, especially at the early time points. In brief, UMAPs for all cell types in stage E6.5 and E6.75 were calculated and force-directed graphs were constructed in line with the original study. Neighbors of cells in trajectory 1 were identified by measuring Euclidean distance to the centroid and cell type annotation (epiblast and primitive steak). With a distance cut-off of 2.6, we imputed 125 ‘similar’ cells at E6.5 and 23 at E6.75. As a result, we have 3,588 cells in trajectory 1 and 3,072 cells in trajectory 2. All gene expression values are normalized to sequencing size factor and log2 transformed. Data handling was performed in R (version 3.6.0).

### 4.2 Macroscale modeling of gene regulatory network

Variance in gene expression was modeled using *modelGeneVar* function in the R package *scran*, with loess span of 0.05 and block settings on 3 sequencing batches. Genes with significantly higher variance than the fitted trend (Benjamini-Hochberg corrected P < 0.05) and positive biological component were selected. Genes were excluded if the mean log2 normalized count is below 10^−3^, on the Y chromosome, or is Xist.

### 4.3 Selection of highly variable genes

Functions of genes and gene products are annotated based on Gene Ontology (GO), a literature-curated reference database (annotation data was downloaded from Mouse Genome Informatics: http://www.informatics.jax.org). In the partially hierarchical structure of the GO directed acyclic graph, child-parent relation links terms of lower-level biological subsystems, such as small complexes or single reactions, to those pertaining to upper-level biological systems, such as organelles or broad processes (Explorable on Mouse Genome Informatics: http://www.informatics.jax.org/vocab/geneontology). To build a macroscale representation of the mouse gene network, we utilized such membership information from the cut-down version of the GO maintained by Mouse Genome Informatics (Mouse GO slim, file name ‘goslimmouse’). This was performed using OWLTools (https://github.com/owlcollab/owltools/wiki/Map2Slim), to aggregate all the lower-level annotations to 14 top-level modules (macroscale functional domains of biological processes). Each annotated gene is tagged with at least one of the 14 biological processes.

### 4.4 Data aggregation

For each trajectory, log2 normalized read counts of all sampled HVGs in each single cell were aggregated at each of the 9 time points (Embryonic day E6.5-E8.5, sampled every 6 hours). For each time points, cells were randomly split into four equal-length sets (four-fold learning scheme). For each set, the average expression level of each gene at a time point was calculated by taking the arithmetic mean of the log2 normalized read counts of the gene in all cells staged at the specific time point. Second, the average expression level of all genes tagged in the same macroscale process were summed at each time point, generating a time-series of state values, representing the gross expression level of each biological process at each time point. As a result, we have 8 sets of data: two trajectories, each consists of four sets of 14 time-series. Each time-series represents the dynamic state values of the macroscale biological process of interest.

### 4.5 Network learning scheme

To construct a causal model of the macroscale network capturing the system dynamics, we built a FSNN (Feedback System Neural Network) (Schoenberg, 2019), which is an ordinary differential equation (ODE)- based machine learning method (1):

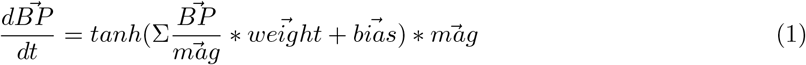

Our FSNN had 14 states 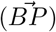 and was initialized with all weights and biases set to 0, thus initially representing a fully disconnected network. The initial value of each state was loaded from the data at E6.5. The FSNN for the model of both trajectories used a single perceptron to model the change in each biological process. The input to each perceptron was the current value of all of the 14 biological processes. The FSNN differential equation was simulated using the RK4 algorithm with a Δt of 1/16 (unit: day). The state values were lineally rescaled using a constant magnitude 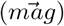 which was calculated as 1.5 times the initial value of the state from the first training set to avoid signal attenuation issues due to the tanh activation function. We trained the FSNN using ¾ of the data and tested on the remaining ¼. The FSNN was constructed and trained using Stella Architect 2.0 which uses Powell’s BOBYQA optimization algorithm as implemented in DLIB version 19.7 to determine the weights and biases. The two trajectories were trained and tested separately, thus generating two distinct models.

### 4.6 Inference on pairwise causal relationship

The two causal models (FSNNs) of the macroscale network were then analyzed to infer direct pairwise impact relations from source process to target process. Here, we utilized the link score measure (2) (Schoenberg et al., 2020) to determine the contribution of one biological process to the change of another in the generated models:

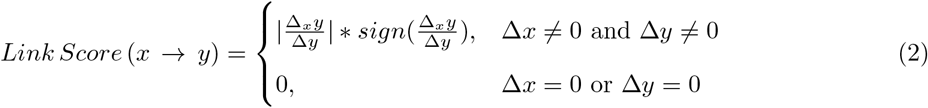

The sign of the link score indicates the polarity of the relationship. A positive value means that the change in the source variable produces a change in the same direction in the target variable, independent of state changes in other variable; while a negative value means the opposite. The magnitude (after normalization across all incoming variables to the target) demonstrates the relative strength of the causal relationship of interest relative to all the other incoming sources to a given target. The mean and variance of link scores were derived from the training and testing sets of time-series data. This analysis was conducted using Stella Architect 2.0.

### 4.7 Feedback loop dominance analysis

To discover the origins of behavior for each causal model from a feedback loop perspective, feedback loop dominance analysis was done using the Loops that Matter method (Schoenberg et al., 2020) implemented in Stella Architect 2.0. The feedback loop discovery process was done using the SPA algorithm because it was impossible to enumerate and measure the 14 factorial feedback loops produced by each FSNN (Eberlein & Schoenberg, 2020). The analysis was performed on just the testing set for each model. The feedback loop dominance analysis measures the share of model behavior which is attributable to each feedback loop in the model at each time point. Those values were averaged over the full simulation period to measure the total contribution of each feedback loop to the behavior of each model over the full simulation time period (Schoenberg & Eberlein, 2020) to clarify the most impactful feedback processes during organogenesis.

### 4.8 Data visualization

A static view of the feedback structure of each model was produced using the link score data from the testing set of each model. Network visualization were created for each model where each node represented a biological process, and each edge represented the average relative link score measured across the simulation period. The number of edges was filtered, and we selected the upper quartile of strongest connections. Gephi (v0.91; Bastian, Heymann, & Jacomy, 2009) was used to create map of the feedback structure. In addition, overtime videos of the progression of causal influences were created using the link scores measured at each Δt of the FSNN, where only the upper quartile of edges were plotted.

## Supplementary Materials

**Supplementary Video 1**. Instantaneous view of direct pairwise causal feedback relationships in the macroscale network model (movie for T1; related to Fig 3).

**Supplementary Video 2**. Instantaneous view of direct pairwise causal feedback relationships in the macroscale network model (movie for T2; related to Fig 3).

**Supplementary Figure 1**. Full matrix of inferred pairwise causal relationships measured by link score (T1; related to Fig 4).

**Supplementary Figure 2**. Full matrix of inferred pairwise causal relationships measured by link score (T2; related to Fig 4).

**Supplementary Figure 3**. Topological measures of the macroscale network models. (A, C) Degree distribution and eccentricity distribution for the macroscale network model (T1). (B, D) Degree distribution and eccentricity distribution for the macroscale network model (T2).

**Supplementary Table 1**. Full Gene Ontology descriptions of the 14 core biological processes.

**Supplementary Table 2**. List of HVGs tagged in each core biological process.

**Supplementary Table 3**. Outcome of model parametrization for the two trajectories.

**Supplementary Table 4**. Outcome of feedback loop dominance analysis (T1).

**Supplementary Table 5**. Outcome of feedback loop dominance analysis (T2).

## References

Ahmadian, M., Suh, J. M., Hah, N., Liddle, C., Atkins, A. R., Downes, M., & Evans, R. M. (2013, May). PPAR signaling and metabolism: the good, the bad and the future. Nature Medicine, 19 (5), 557–566. doi:10.1038/nm.3159

Aibar, S., González-Blas, C. B., Moerman, T., Huynh-Thu, V. A., Imrichova, H., Hulselmans, G., … Aerts, S. (2017, November). SCENIC: single-cell regulatory network inference and clustering. Nature Methods, 14 (11), 1083–1086. doi:10.1038/nmeth.4463

Arnold, S. J., & Robertson, E. J. (2009, February). Making a commitment: cell lineage allocation and axis patterning in the early mouse embryo. Nature Reviews Molecular Cell Biology, 10 (2), 91–103. Retrieved 2024-02-01, from https://www.nature.com/articles/nrm2618 doi:10.1038/nrm2618

Barresi, M. J. F., & Gilbert, S. F. (2024). Developmental biology (13. edition ed.). New York, NY: Oxford University Press.

Basson, M. A. (2012, June). Signaling in cell differentiation and morphogenesis. Cold Spring Harbor Perspectives in Biology, 4 (6), a008151. doi:10.1101/cshperspect.a008151

Bastian, M., Heymann, S., & Jacomy, M. (2009, March). Gephi: An Open Source Software for Exploring and Manipulating Networks. Proceedings of the International AAAI Conference on Web and Social Media, 3 (1), 361–362. Retrieved 2024-02-01, from https://ojs.aaai.org/index.php/ICWSM/article/view/13937 doi:10.1609/icwsm.v3i1.13937

Bult, C. J., Blake, J. A., Smith, C. L., Kadin, J. A., Richardson, J. E., & Mouse Genome Database Group. (2019, January). Mouse Genome Database (MGD) 2019. Nucleic Acids Research, 47 (D1), D801–D806. doi:10.1093/nar/gky1056

Cao, J., Spielmann, M., Qiu, X., Huang, X., Ibrahim, D. M., Hill, A. J., … Shendure, J. (2019, February). The single-cell transcriptional landscape of mammalian organogenesis. Nature, 566 (7745), 496–502. doi:10.1038/s41586-019-0969-x

Casadiego, J., Nitzan, M., Hallerberg, S., & Timme, M. (2017, December). Model-free inference of direct network interactions from nonlinear collective dynamics. Nature Communications, 8 (1), 2192. doi:10.1038/s41467-017-02288-4

Causton, H. C. (2020, January). Metabolic rhythms: A framework for coordinating cellular function. The European Journal of Neuroscience, 51 (1), 1–12. doi:10.1111/ejn.14296

Cheng, S., Pei, Y., He, L., Peng, G., Reinius, B., Tam, P. P. L., … Deng, Q. (2019, March). Single-Cell RNA-Seq Reveals Cellular Heterogeneity of Pluripotency Transition and X Chromosome Dynamics during Early Mouse Development. Cell Reports, 26 (10), 2593–2607.e3. doi:10.1016/j.celrep.2019.02.031

Davidson, E. H., Rast, J. P., Oliveri, P., Ransick, A., Calestani, C., Yuh, C.-H., … Bolouri, H. (2002, March). A genomic regulatory network for development. Science (New York, N.Y.), 295 (5560), 1669–1678. doi:10.1126/science.1069883

Eberlein, R., & Schoenberg, W. (2020, May). Finding the Loops that Matter. arXiv. Retrieved 2024-02-01, from http://arxiv.org/abs/2006.08425 (2006.08425 [cs])

Ferrell, J. E. (2012, June). Bistability, bifurcations, and Waddington’s epigenetic landscape. Current biology: CB, 22 (11), R458–466. doi:10.1016/j.cub.2012.03.045

Ferrell, J. E. (2016, February). Perfect and Near-Perfect Adaptation in Cell Signaling. Cell Systems, 2 (2), 62–67. doi:10.1016/j.cels.2016.02.006

Fondi, M., & Liò, P. (2015). Genome-scale metabolic network reconstruction. Methods in Molecular Biology (Clifton, N.J.), 1231, 233–256. doi:10.1007/978-1-4939-1720-415

Fraser, H. B., Hirsh, A. E., Steinmetz, L. M., Scharfe, C., & Feldman, M. W. (2002, April). Evolutionary rate in the protein interaction network. Science (New York, N.Y.), 296 (5568), 750–752. doi:10.1126/science.1068696

Fujii, K., Takeishi, N., Hojo, M., Inaba, Y., & Kawahara, Y. (2020, February). Physically-interpretable classification of biological network dynamics for complex collective motions. Scientific Reports, 10 (1), 3005. doi:10.1038/s41598-020-58064-w

Gomes, A. P., & Blenis, J. (2015, August). A nexus for cellular homeostasis: the interplay between metabolic and signal transduction pathways. Current Opinion in Biotechnology, 34, 110–117. doi:10.1016/j.copbio.2014.12.007

Goodsell, D. S., Olson, A. J., & Forli, S. (2020, June). Art and Science of the Cellular Mesoscale. Trends in Biochemical Sciences, 45 (6), 472–483. doi:10.1016/j.tibs.2020.02.010

Groß, A., Kracher, B., Kraus, J. M., Kühlwein, S. D., Pfister, A. S., Wiese, S., … Kestler, H. A. (2019). Representing dynamic biological networks with multi-scale probabilistic models. Communications Biology, 2, 21. doi:10.1038/s42003-018-0268-3

Han, J.-D. J., Bertin, N., Hao, T., Goldberg, D. S., Berriz, G. F., Zhang, L. V., … Vidal, M. (2004, July). Evidence for dynamically organized modularity in the yeast protein-protein interaction network. Nature, 430 (6995), 88–93. doi:10.1038/nature02555

Harush, U., & Barzel, B. (2017, December). Dynamic patterns of information flow in complex networks. Nature Communications, 8 (1), 2181. doi:10.1038/s41467-017-01916-3

Jeong, H., Tombor, B., Albert, R., Oltvai, Z. N., & Barabási, A. L. (2000, October). The large-scale organization of metabolic networks. Nature, 407 (6804), 651–654. doi:10.1038/35036627

Kang, J., Benjamin, D. I., Kim, S., Salvi, J. S., Dhaliwal, G., Lam, R., … Rando, T. A. (2024, January). Depletion of SAM leading to loss of heterochromatin drives muscle stem cell ageing. Nature Metabolism, 6 (1), 153–168. Retrieved 2024-02-01, from https://www.nature.com/articles/s42255-023-00955-z doi:10.1038/s42255-023-00955-z

Kurz, F. T., Kembro, J. M., Flesia, A. G., Armoundas, A. A., Cortassa, S., Aon, M. A., & Lloyd, D. (2017, January). Network dynamics: quantitative analysis of complex behavior in metabolism, organelles, and cells, from experiments to models and back. Wiley Interdisciplinary Reviews. Systems Biology and Medicine, 9 (1). doi:10.1002/wsbm.1352

Liu, Y., Colby, J. K., Zuo, X., Jaoude, J., Wei, D., & Shureiqi, I. (2018, October). The Role of PPAR-in Metabolism, Inflammation, and Cancer: Many Characters of a Critical Transcription Factor. International Journal of Molecular Sciences, 19 (11), 3339. doi:10.3390/ijms19113339

Lodhi, I. J., & Semenkovich, C. F. (2014, March). Peroxisomes: a nexus for lipid metabolism and cellular signaling. Cell Metabolism, 19 (3), 380–392. doi:10.1016/j.cmet.2014.01.002

Lu, R., Markowetz, F., Unwin, R. D., Leek, J. T., Airoldi, E. M., MacArthur, B. D., … Lemischka, I. R. (2009, November). Systems-level dynamic analyses of fate change in murine embryonic stem cells. Nature, 462 (7271), 358–362. doi:10.1038/nature08575

Luscombe, N. M., Babu, M. M., Yu, H., Snyder, M., Teichmann, S. A., & Gerstein, M. (2004, September). Genomic analysis of regulatory network dynamics reveals large topological changes. Nature, 431 (7006), 308–312. doi:10.1038/nature02782

Ma, J., Yu, M. K., Fong, S., Ono, K., Sage, E., Demchak, B., … Ideker, T. (2018, April). Using deep learning to model the hierarchical structure and function of a cell. Nature Methods, 15 (4), 290–298. doi:10.1038/nmeth.4627

Macarthur, B. D., Ma’ayan, A., & Lemischka, I. R. (2009, October). Systems biology of stem cell fate and cellular reprogramming. Nature Reviews. Molecular Cell Biology, 10 (10), 672–681. doi:10.1038/nrm2766

Merkuri, F., Rothstein, M., & Simoes-Costa, M. (2024, January). Histone lactylation couples cellular metabolism with developmental gene regulatory networks. Nature Communications, 15 (1), 90. Retrieved 2024-02-01, from https://www.nature.com/articles/s41467-023-44121x-1 doi:10.1038/s41467-023-44121-1

Neph, S., Stergachis, A. B., Reynolds, A., Sandstrom, R., Borenstein, E., & Stamatoyannopoulos, J. A. (2012, September). Circuitry and dynamics of human transcription factor regulatory networks. Cell, 150 (6), 1274–1286. doi:10.1016/j.cell.2012.04.040

Pan, G., Li, J., Zhou, Y., Zheng, H., & Pei, D. (2006, August). A negative feedback loop of transcription factors that controls stem cell pluripotency and self-renewal. FASEB journal: official publication of the Federation of American Societies for Experimental Biology, 20 (10), 1730–1732. doi:10.1096/fj.05-5543fje

Papatsenko, D., Waghray, A., & Lemischka, I. R. (2018, May). Feedback control of pluripotency in embryonic stem cells: Signaling, transcription and epigenetics. Stem Cell Research, 29, 180–188. doi:10.1016/j.scr.2018.02.012

Peixoto, T. P., & Rosvall, M. (2017, September). Modelling sequences and temporal networks with dynamic community structures. Nature Communications, 8 (1), 582. doi:10.1038/s41467-017-00148-9

Pijuan-Sala, B., Griffiths, J. A., Guibentif, C., Hiscock, T. W., Jawaid, W., Calero-Nieto, F. J., … Göttgens, B. (2019, February). A single-cell molecular map of mouse gastrulation and early organogenesis. Nature, 566 (7745), 490–495. doi:10.1038/s41586-019-0933-9

Rockne, R. C., Branciamore, S., Qi, J., Frankhouser, D. E., O’Meally, D., Hua, W.-K., … Marcucci, G. (2020, August). State-Transition Analysis of Time-Sequential Gene Expression Identifies Critical Points That Predict Development of Acute Myeloid Leukemia. Cancer Research, 80 (15), 3157–3169. doi:10.1158/0008-5472.CAN-20-0354

Rozenblatt-Rosen, O., Regev, A., Oberdoerffer, P., Nawy, T., Hupalowska, A., Rood, J. E., … Human Tumor Atlas Network (2020, April). The Human Tumor Atlas Network: Charting Tumor Transitions across Space and Time at Single-Cell Resolution. Cell, 181 (2), 236–249. doi:10.1016/j.cell.2020.03.053

Schoenberg, W. (2019). Feedback System Neural Networks for Inferring Causality in Directed Cyclic Graphs. Retrieved 2024-02-01, from https://arxiv.org/abs/1908.10336 (Publisher: arXiv Version Number: 2) doi:10.48550/ARXIV.1908.10336

Schoenberg, W., Davidsen, P., & Eberlein, R. (2020, May). Understanding model behavior using loops that matter. arXiv. Retrieved 2024-02-01, from http://arxiv.org/abs/1908.11434 (1908.11434 [physics])

Schoenberg, W., & Eberlein, R. (2020, May). Seamlessly Integrating Loops That Matter into Model Development and Analysis. arXiv. Retrieved 2024-02-01, from http://arxiv.org/abs/2005.14545 (2005.14545 [cs, math])

Shin, S., Venturelli, O. S., & Zavala, V. M. (2019, March). Scalable nonlinear programming framework for parameter estimation in dynamic biological system models. PLoS computational biology, 15 (3), e1006828. doi:10.1371/journal.pcbi.1006828

Sladitschek, H. L., Fiuza, U.-M., Pavlinic, D., Benes, V., Hufnagel, L., & Neveu, P. A. (2020, May). MorphoSeq: Full Single-Cell Transcriptome Dynamics Up to Gastrulation in a Chordate. Cell, 181 (4), 922–935.e21. doi:10.1016/j.cell.2020.03.055

Tam, P. P. L., & Loebel, D. A. F. (2007, May). Gene function in mouse embryogenesis: get set for gastrulation. Nature Reviews. Genetics, 8 (5), 368–381. doi:10.1038/nrg2084

The Gene Ontology Consortium. (2017, January). Expansion of the Gene Ontology knowledgebase and resources. Nucleic Acids Research, 45 (D1), D331–D338. doi:10.1093/nar/gkw1108

Tu, B. P., Kudlicki, A., Rowicka, M., & McKnight, S. L. (2005, November). Logic of the yeast metabolic cycle: temporal compartmentalization of cellular processes. Science (New York, N.Y.), 310 (5751), 1152–1158. doi:10.1126/science.1120499

Wellen, K. E., & Thompson, C. B. (2012, March). A two-way street: reciprocal regulation of metabolism and signalling. Nature Reviews. Molecular Cell Biology, 13 (4), 270–276. doi:10.1038/nrm3305

Xiong, L., & Garfinkel, A. (2022, October). A common pathway to cancer: Oncogenic mutations abolish p53 oscillations. Progress in Biophysics and Molecular Biology, 174, 28–40. Retrieved 2022-09-02, from https://linkinghub.elsevier.com/retrieve/pii/S0079610722000633 doi:10.1016/j.pbiomolbio.2022.06.002

Xiong, L. I., & Garfinkel, A. (2023, August). Are physiological oscillations physiological ? The Journal of Physiology, JP285015. Retrieved 2023-11-07, from https://physoc.onlinelibrary.wiley.com/doi/10.1113/JP285015 doi:10.1113/JP285015

Young, R. A. (2011, March). Control of the embryonic stem cell state. Cell, 144 (6), 940–954. doi:10.1016/j.cell.2011.01.032

Zou, Y., Donner, R. V., Marwan, N., Donges, J. F., & Kurths, J. (2019, January). Complex network approaches to nonlinear time series analysis. Physics Reports, 787, 1–97. Retrieved 2024-02-01, from https://linkinghub.elsevier.com/retrieve/pii/S037015731830276X doi:10.1016/j.physrep.2018.10.005

