## Supplementary figures and images for "Learning feedback loops in transcriptome network during cell fate transition"

### Fig_S1.pdf

#### Endoderm set1_aug HL0 - Causality Matrix

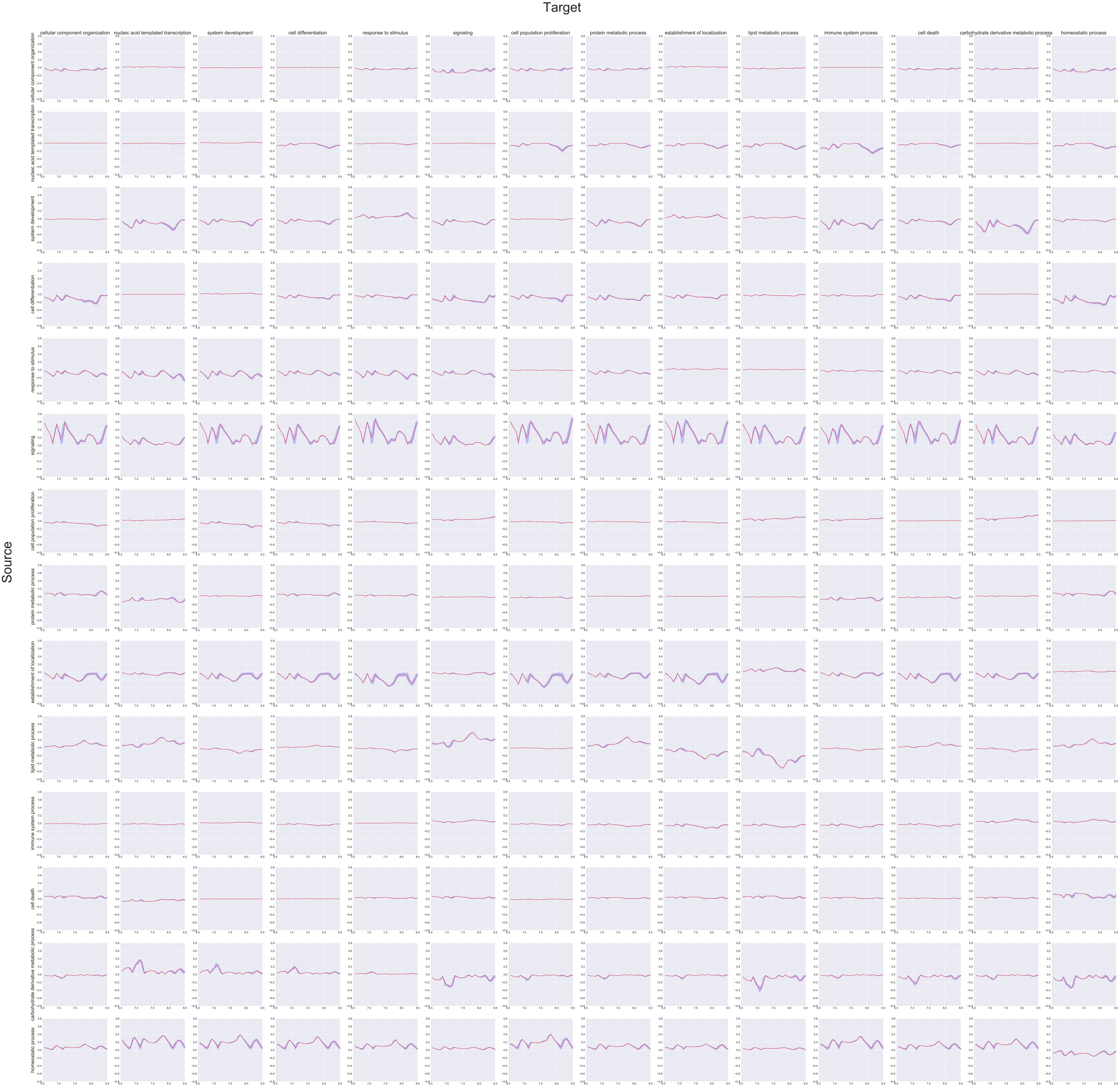

### Fig_S2.pdf

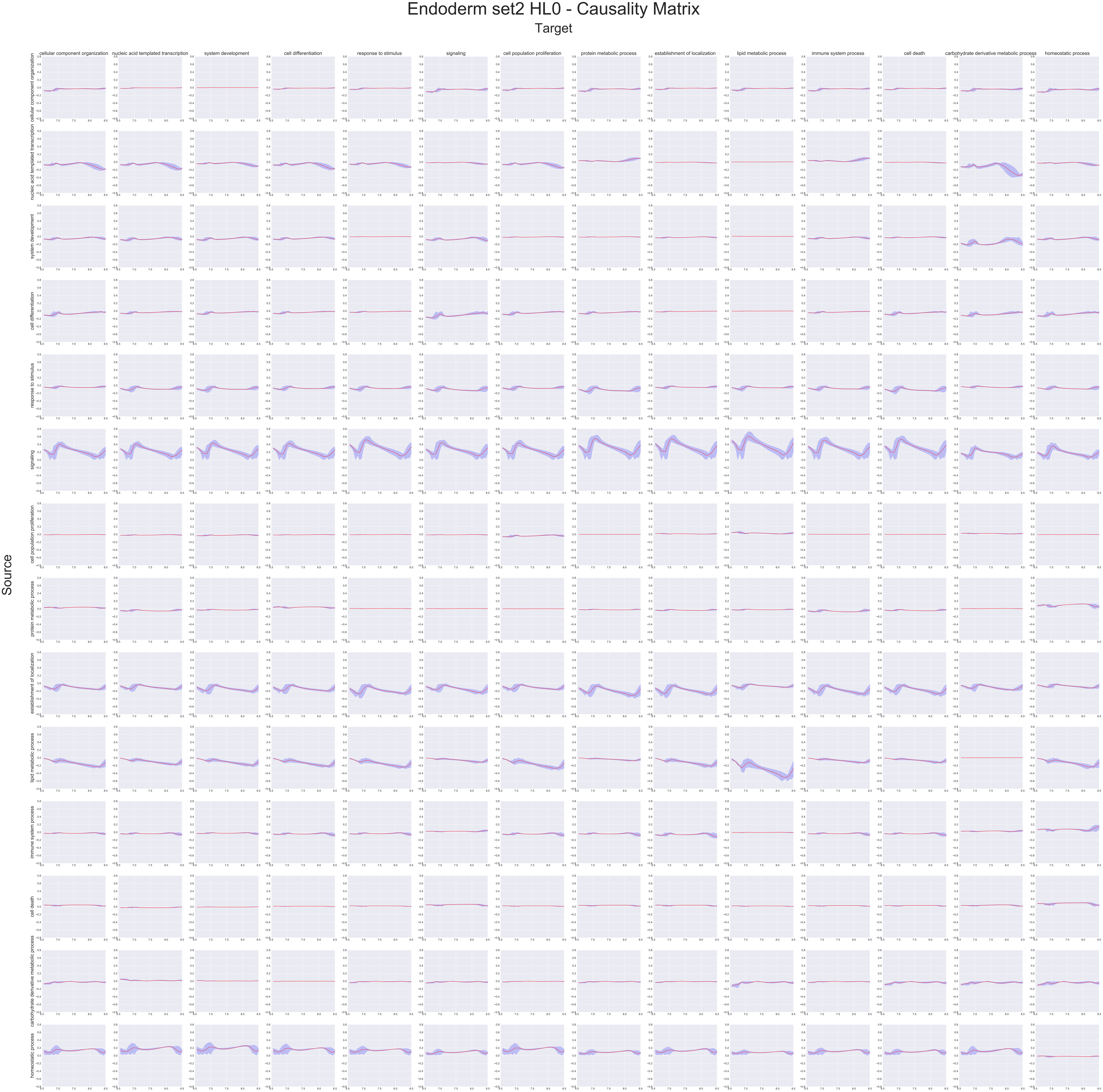

### Fig_S3.pdf

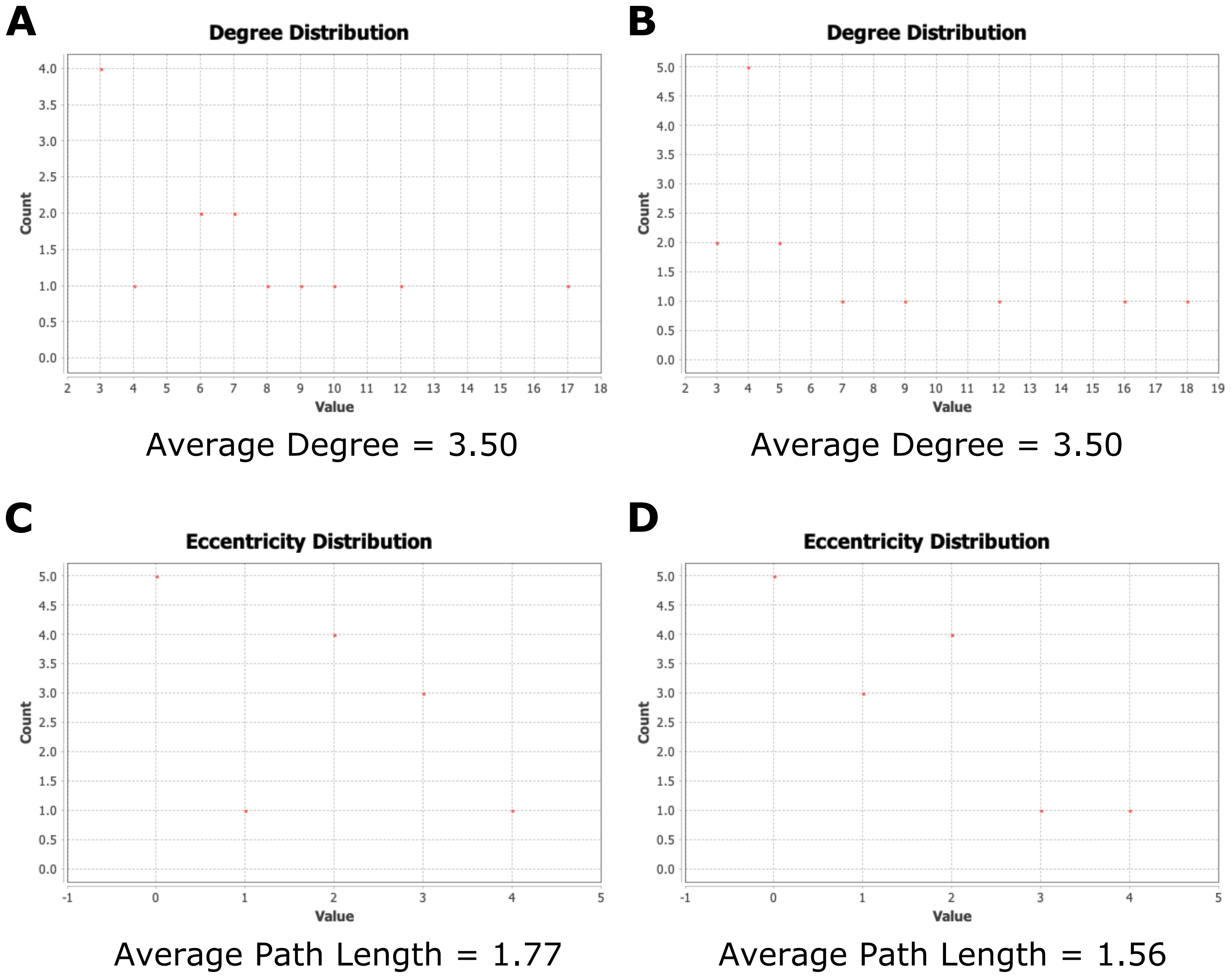
